# Lack of canonical PAR4 activation is associated with reduced arterial and venous thrombosis in mice

**DOI:** 10.64898/2026.07.20.739624

**Authors:** Robert H. Lee, Jenna R. Severa, David S. Paul, Woosuk S. Hur, Swati Sharma, Dale O. Cowley, Matthew J. Flick, Wolfgang Bergmeier, Nigel Mackman, Timothy J. Stalker, Silvio Antoniak

## Abstract

**Background:** Protease-activated receptor 4 (PAR4) is the only functional thrombin receptor on mouse platelets. Expression and activation of platelet PAR4 was shown to influence hemostatic plug stability and contributes to thrombosis in different murine arterial and venous thrombosis models. PAR4 activation by thrombin and other serine proteases occurs at the canonical activation site at Arginine (Arg) 59 in mice and Arg47 in humans. If murine PAR4 has functional non-canonical activation sites as shown for other PARs is unknown.

**Objective:** To investigate canonical and potentially non-canonical PAR4 signaling in mice, we generated a mouse model expressing a functional, thrombin-cleavage resistant PAR4 by changing Arg59 to Alanine (Ala) 59 in murine PAR4 (PAR4^R59A^). PAR4^R59A^ mice were used to assess the impact of impaired canonical (thrombin)-dependent PAR4 signaling on hemostasis and thrombosis in mice.

**Methods:** We analyzed platelet aggregation, platelet integrin activation and α-granule release *ex vivo*. Hemostasis and thrombosis in PAR4^R59A^ and their control mice was compared using the jugular vein needle puncture injury-induced hemostasis model, and the ferric chloride-induced carotid artery and electrolytic injury-induced femoral vein thrombosis models.

**Results:** Platelets of PAR4^R59A^ mice did not response to thrombin but responded normally to PAR4 agonist peptide (PAR4AP) stimulation in aggregation assays. Platelets of PAR4^R59A^ and their control mice exhibited comparable responses to ADP, convulxin or PAR4AP regarding integrin activation and α-granular release. PAR4^R59A^ mice exhibited impaired hemostasis in the jugular vein needle puncture model, and were protected from ferric chloride-induced arterial thrombosis and from electrolytic injury induced thrombosis in the femoral vein.

**Conclusion:** The novel PAR4^R59A^ mouse expresses a thrombin-insensitive but still functional PAR4. We propose that the new mouse line will increase the *in vivo* investigation of canonical PAR4 signaling pathways and may reveal unknown non-canonical PAR4 signaling in different pathologies.

## Introduction

Protease-activated receptors (PARs) are membrane receptors that detect changes in the proteolytic milieu of the extracellular compartment. Serine proteases are the most common activating proteases for PARs^1^. There are 4 PARs identified. PAR1, also known as factor 2 receptor (F2R) or thrombin receptor (TR), was first described as the receptor responsible for thrombin-dependent activation of human platelets^2^. PAR4 (F2R-like 3, F2RL3) is the other known functional TR on human platelets^3^. In most other species including mice, PAR4 is the only functional TR on platelets^3^. This fact underscores the biological importance of PAR4 for platelet activation. Thrombin cleaves human PAR4 at Arg47 (R47) and murine PAR4 at Arg59 (R59). Cleavages at these sites are termed “canonical cleavage” which leads to strong cellular responses including platelet activation. Besides thrombin, canonical PAR4 activation was also reported for the potent blood-borne protease plasmin^4, 5^. In addition, trypsin and immune cell associated proteases such as cathepsin G (CatG) can also activate PAR4^6–10^. If murine PAR4 has any functional non-canonical activation sites is unknown. Recently, we presented the generation of mice expressing a conditional PAR4 gene (PAR4^fl/fl^)^11^, which allows a cell-specific investigation of PAR4 *in vivo*. Using Pf4-Cre to delete PAR4 in megakaryocytes/platelets, we showed that platelet PAR4 is needed for stable hemostatic plug formation in a laser-induced vascular injury model^11^. Additionally, platelet PAR4 plays a role in arterial and venous thrombosis^11^.

Until now, an investigation of global PAR4 signaling deficiency in mice was only possible by using mice carrying a disrupted PAR4 gene (PAR4^-/-^)^12^. The PAR4 gene disruption in the PAR4^-/-^ mice was achieved by introducing a β-galactosidase (LacZ) coding DNA before the signal peptide encoding sequence of murine PAR4^12^. Platelets from these transgenic PAR4^-/-^ mice are unresponsive to thrombin or PAR4AP stimulation^12^.

To investigate the lack of canonical PAR4 signaling *in vivo*, we generated mice carrying an Arg59 to Ala59 mutation in the canonical cleavage site of murine PAR4 (PAR4^R59A^). The R59A mutation in murine PAR4 results in platelets that are insensitive to thrombin yet retain otherwise normal responsiveness. Compared to wild-type mice, PAR4^R59A^ mice exhibit impaired hemostasis in a jugular vein needle puncture injury model. More importantly, PAR4^R59A^ mice showed protection in the ferric chloride-induced carotid artery thrombosis model and exhibit reduced platelet accumulation in the femoral vein electrolytic injury model. The PAR4^R59A^ mouse line provides a new tool and the opportunity to investigate a selective lack of canonical PAR4 signaling and may reveal non-canonical PAR4 signaling *in vivo*.

## Methods

### Generation of an F2rl3 (PAR4) R59A knock-in mouse by CRISPR/Cas9-mediated genome editing

#### CRISPR/Cas9 Reagents

Cas9 guide RNAs (gRNAs) targeting the F2rl3 R59 codon in exon 2 gene were identified using Benchling software (benchling.com) and analyzed for on-target and off-target profiles using CRISPOR (crispor.tefor.net). Three gRNAs were selected for activity testing. Guide RNAs were cloned into a T7 promoter vector (UNC Animal Models Core) followed by in vitro transcription (HiScribe T7 High Yield RNA Synthesis Kit, New England BioLabs, Ipswich, MA) and RNeasy spin column purification (QIAGEN, Germantown, MD). Functional testing was performed by transfecting an immortalized mouse embryonic fibroblast cell line (UNC Animal Models Core) with gRNA and recombinant Cas9 protein (UNC Animal Models Core/UNC Protein Purification Core Facility). The target region was amplified from transfected cell lysates using primers listed below and analyzed by Sanger sequencing followed by ICE Analysis (Synthego). Two guide RNAs were selected for genome editing in embryos: F2rl3-g92B (protospacer sequence 5’-ATTTGCCCGGGTAGCCTCG-3’) and F2rl3-g83T (protospacer sequence 5’-GCCTAATCCACGAGGCTACC-3’). A donor oligonucleotide (Sigma-Aldrich, St. Louis, MO), was designed to facilitate homologous recombination to introduce the R59A (CGA to GCT) mutation. The donor sequence was 5’-CTGGGTCCCACAGTAGAACTCAAGGAGCCGAAGTCCTCAGACAAGCCTAATCCAgctGGCTACCCGGGCAAATTCTGTGCCAACGACAGTGACACGCTGGAGCTCCCGGC - 3’.

### Embryo Electroporation

C57BL/6J zygotes were electroporated with one of two CRISPR reagent mixes (one with gRNA g92B and one with gRNA g83T). Each mix was comprised of 600 nM recombinant Cas9 protein, 24 ng/μl gRNA and 400 ng/μl donor oligonucleotide in Opti-MEM I (Thermo Fisher Scientific, Waltham, MA). Electroporated embryos were implanted in pseudopregnant recipient females and resulting pups were screened by PCR and sequencing for the presence of the R59A allele. Positive founders were identified from both electroporation mixes, with higher positive founder rates from the g92B gRNA mix. Two male founders harboring the R59A mutation from the gRNA g92B electroporation mix were mated to wild-type C57BL/6J females to establish colonies with the R59A allele.

### Genotyping

Animals were genotyped by PCR amplification of the target region with primers F2rl3-ScF1 (5’-CCATTAACTGAGTGCTGGAGTGG −3’) and F2rl3-ScR1 (5’-AGCCAGTCGTGGTGGCAG −3’). The resulting 460 bp product was sequenced with primer F2rl3-SqF1 (5’-ACACTGAGGCACCCACAATG −3’). The R59A mutation introduced a *PvuII* digestion site in the PAR4 coding sequence. The 460 bp PCR product was digested with *PvuII* for differential digestion of the R59A and WT allele PCR products.

### Platelet function analysis

Venous blood was collected into low molecular weight heparin (Lovenox, Sanofi, France)^11^. For aggregometry studies, washed platelets were resuspended in modified Tyrode’s buffer at a final concentration of 2.5 × 10^8^ platelets/mL in a Chrono-log 4-channel optical aggregation system (Chrono-log, Havertown, PA) and stimulated with α-thrombin (1 U/ml, Enzyme Research Laboratories, South Bend, IN), PAR4AP (250 µM, GL Biochem, Shanghai, China) or convulxin (Cvx, 100 ng/ml, purchased from Kenneth Clemetson, Theodor Kocher Institute, University of Berne, Switzerland). To analyze αIIbβ3 integrin activation or α-granular release, binding of JON/A PE (Emfret Analytics) or anti-CD62P-AlexaFluor488 (clone RB40.34, BD Bioscience) to resting, ADP (5 μM), Cvx (30 or 200 ng/mL) or PAR4AP (150 or 400μM) stimulated platelets was quantified by flow cytometry^11^.

### Jugular vein needle puncture model

Male and female C57BL6/J and homozygous PAR4^R59A^ mice were between the age of 8-16 weeks with a weight less than 30 g. The jugular vein needle puncture injury model was performed essentially as previously described^13, 14^. Anti-GPIbβ 650 (Emfret, X650), Alexa 568-labelled anti-P-selectin (BD Biosciences, clone RB40.34), and Alexa 488 labelled fibrinogen (Invitrogen) were infused via retroorbital injection immediately prior to surgery. The right external jugular vein was exposed, cleaned of connective tissue, and pierced with a 200 µm diameter needle. At 5 minutes post-injury, the mouse was perfused with phosphate buffered saline, followed by fixation with 4% paraformaldehyde and excised. Excised veins were opened, pinned flat in a silicone bottomed dish, and submerged in fixative. Tissue clearing was performed by submerging the tissue in 50% thiodiethanol (TDE) overnight, followed by 90% TDE for 3 hours prior to imaging. Images were acquired using a Nikon A1R confocal microscope with a 25x silicone (1.05 NA) objective. All raw images were processed with Nikon NIS-Elements denoising algorithm and deconvolved (Richardson-Lucy, 10 iterations). Visualization and analyses were completed using Imaris (version 10.2, Oxford Instruments).

### Ferric chloride (FeCl_3_) carotid artery thrombosis

Isoflurane (2-3%) anesthetized male mice were placed in a supine position, and the right carotid artery was isolated by blunt dissection^15^. A 10% FeCl_3_-soaked piece of filter paper (1×2mm) was applied to the ventral surface of the exposed carotid artery for 3 min. Then, a microvascular ultrasonic flow probe (MA0.5PSB, Transonic Systems Inc, Ithaca, NY, USA) was used to measure blood flow for 30 min. Artery occlusion was defined as a cessation of blood flow (0 ml/min) for ≥2 min^15^.

### Electrolytic femoral vein thrombosis

Male mice were subjected to a femoral vein electrolytic injury model of venous thrombosis. Mice were placed, supine, on a warming pad prior to removal of fur from the inner surface of the left hind limb with depilatory cream. A 1-2 cm incision was made in the inguinal region to expose the femoral vein just prior to intravenous infusion of Alexa Fluor 488-labeled anti-GPIX antibody (3.75 μg/mouse, clone Xia.B4, Emfret Analytics), Alexa Fluor 647-labeled anti-fibrin antibody (2 μg/mouse, clone 59D8, in-house) to label platelets and fibrin, respectively. Electrolytic injuries were achieved using a 100 µm diameter stainless steel wire to apply a 1.5V direct current (0.02 Amps) generated by a linear power supply (model: DP832, Rigol Technologies) to the ventral surface of the femoral vein for 30 seconds. Accumulation of platelets and fibrin was monitored by intravital video-microscopy using a stereo microscope (SMZ25, Nikon Corp.) coupled to an ORCA Flash 4.0 digital camera (Hamamatsu, Japan). Data were analyzed using NIS-Elements software (Nikon, Corp.).

All animal experiments were performed in accordance with the guidelines of the animal care and use committees of the University of North Carolina at Chapel Hill and Thomas Jefferson University and complied with National Institutes of Health guidelines.

### Statistics

Statistical analyses were performed with GraphPad Prism 10 (GraphPad Software Inc., La Jolla, CA). If not stated otherwise, data are represented as mean±SD. For 2-group comparison, the Mann-Whitney test was used. Bleeding times were analyzed by log-rank test. A *p* value ≤0.05 was regarded as significant.

## Results

### The R59A mutation in murine PAR4 leads to thrombin insensitive platelets

Platelets from wild-type and homozygous PAR4^R59A^ mice were stimulated with α-thrombin, PAR4AP and Cvx and their response analyzed by light transmission aggregation. Compared to control platelets, platelets from PAR4^R59A^ mice did not aggregate when stimulated with α-thrombin (**Fig. 1A**). However, platelets of both genotypes aggregated when stimulated with PAR4AP (**Fig. 1B**) or Cvx (**Fig. 1C**). Platelet αIIbβ3 integrin activation (**Fig. 1D**) and α-granular release (**Fig. 1E**) was comparable between the two genotypes when stimulated with ADP, Cvx or PAR4AP. The data show that the R59A mutation in murine PAR4 leads to thrombin-insensitive platelets. Importantly, PAR4^R59A^ platelets still express functional PAR4 and have normal responses to ADP, Cvx and PAR4AP.

**Figure 1:**
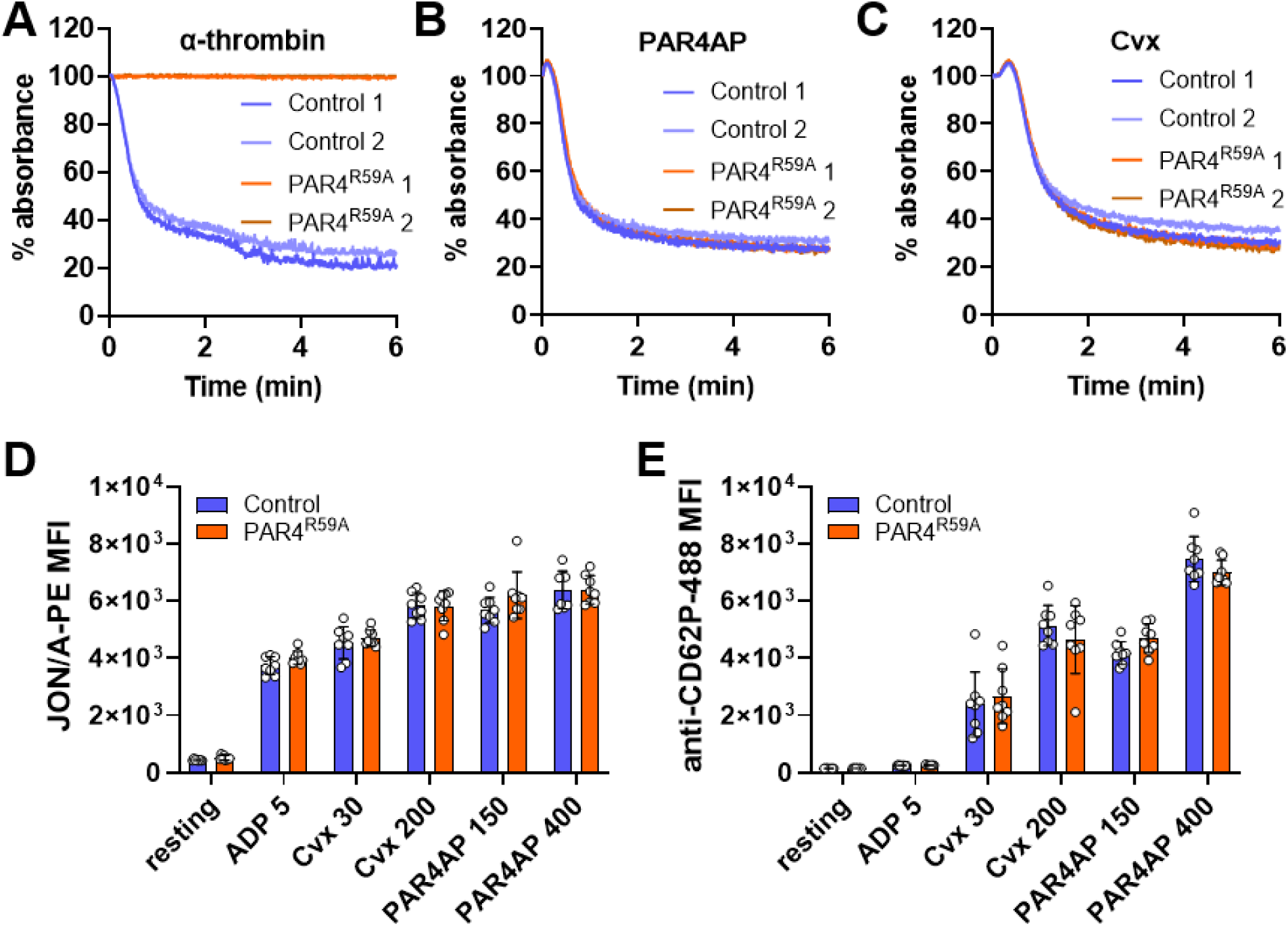
Platelets of PAR4^R59A^ mice lack the response to thrombin. Platelet aggregation of washed control or PAR4^R59A^ mouse platelets stimulated with A) α-thrombin (1 U/ml), B) PAR4 agonist peptide (PAR4AP, 250 μM) or C) convulxin (Cvx, 100 ng/mL). D) αIIbβ3 integrin activation (JON/A PE) and E) α-granular release (anti-CD62P-488, p-Selectin) of platelets in diluted whole blood samples from PAR4^R59A^ and control mice after stimulation with ADP (5μM), Cvx (30 or 200ng/mL) or PAR4AP (150 or 400μM). Mean fluorescence intensity (MFI) was quantified by flow cytometry. Data are shown as mean±SD and analyzed by 2-way ANOVA.

### PAR4^R59A^ mice exhibited impaired hemostasis in the jugular vein needle puncture injury model

We previously reported that mice lacking platelet PAR4 showed signs of impaired hemostatic plug stability in a saphenous vein laser injury model^11^. Here, we used the jugular vein needle puncture injury model to investigate the lack of canonical PAR4 signaling in hemostasis. Mice with the R59A mutation in PAR4 exhibited reduced platelet accumulation quantified by GPIbβ staining of the intraluminal hemostatic plug 5 min after needle injury of the jugular vein (**Fig. 2A**). The hemostatic plug in PAR4^R59A^ mice exhibited reduced signs for platelet activation quantified as CD62P (P-selectin) surface exposure compared to control mice (**Fig. 2B**). The reduced platelet accumulation and CD62P exposure was further associated with reduced fibrin accumulation within the hemostatic plug of PAR4^R59A^ mice compared to control mice (**Fig. 2C**). Together, mice with the PAR4^R59A^ mutation exhibited prolonged bleeding times of the jugular vein compared to control mice (**Fig. 2D**). The data shows that mutation of the canonical activation site of murine PAR4 results in impaired hemostatic plug formation in a needle injury model of the jugular vein.

**Figure 2:**
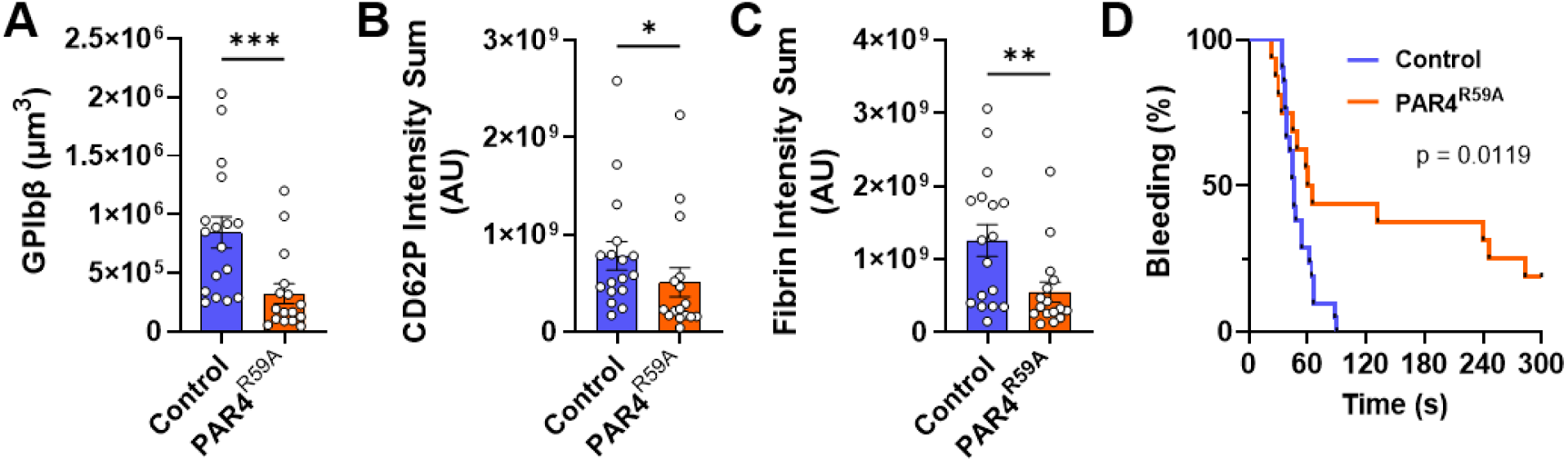
Mice expressing PAR4^R59A^ exhibit impaired hemostasis in the jugular vein needle puncture injury model. PAR4^R59A^ and control mice were subjected to the jugular vein needle puncture injury model. Hemostatic plugs (after 5 mins) were analyzed for A) platelet volume (GPIbβ), B) P-selectin expression (CD62P) and C) fibrin content by confocal microscopy. D) Bleeding time from the injury are shown as Kaplan-Meier plot over the duration of 5 mins. Data are shown as mean±SEM and analyzed by Mann-Whitney test (A-C) or Log-rank (Mantel-Cox) test (D). \**p*<0.05, \*\**p*<0.01 and \*\*\**p*<0.005.

### PAR4^R59A^ mice exhibit reduced arterial thrombosis

Recently, we showed that platelet PAR4 contributes to arterial thrombosis in the murine ferric chloride-induced carotid artery injury model^11^. To test if this was due to canonical PAR4 signaling, we subjected the PAR4^R59A^ and their control mice to the same injury model. While control mice showed carotid artery occlusion, PAR4^R59A^ mice did not occlude over the observation period of up to 30 min post injury (**Fig. 3**). The data shows that PAR4 activation at the canonical site is responsible for occlusive thrombi after ferric chloride-induced carotid artery injury in mice.

**Figure 3:**
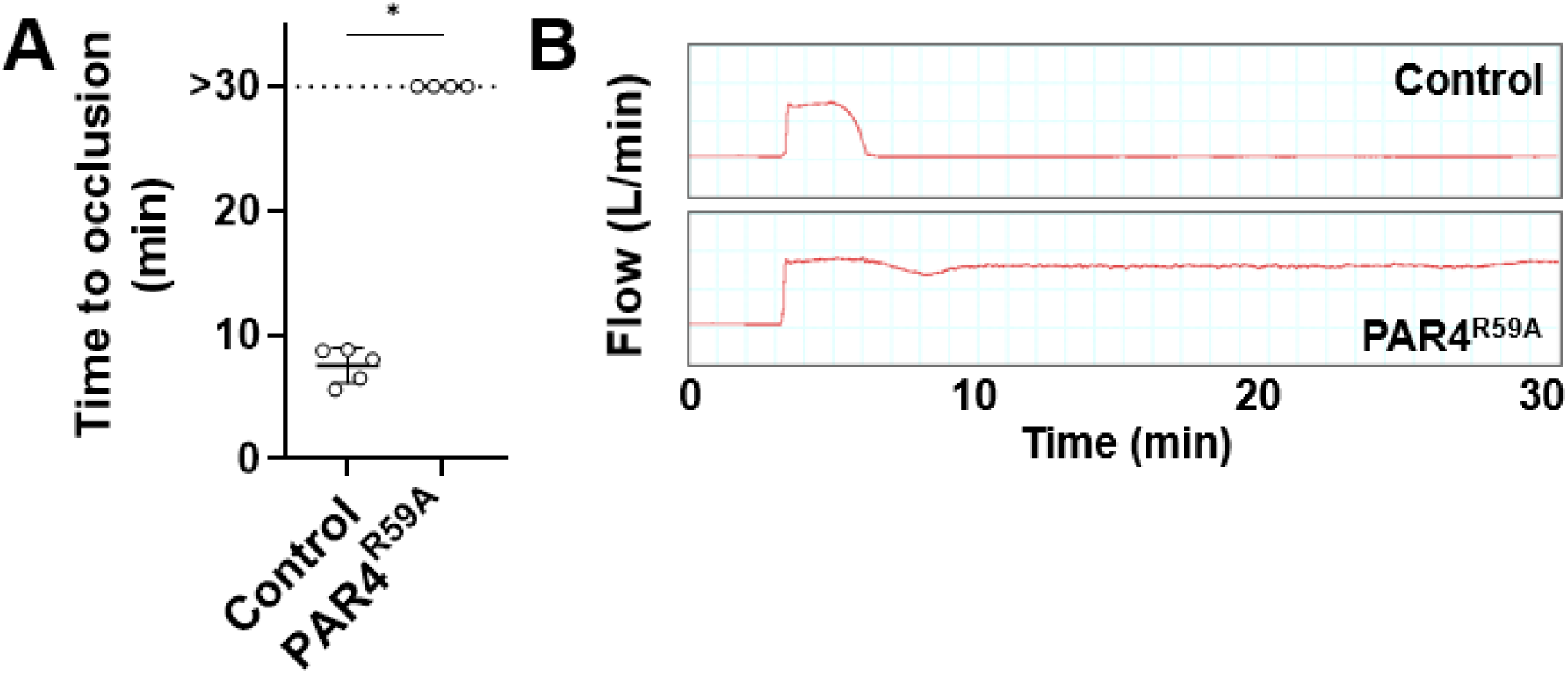
PAR4^R59A^ mice are completely protected from arterial thrombosis. PAR4^R59^ and their control mice were subjected to a ferric chloride-induced carotid artery thrombosis model. A) Occlusion times are shown after 10% FeCl_3_ for 3 min. Occlusion was defined as no blood flow for ≥ 2 min over the duration of 30 min. B) Representative blood flow traces. Data is presented as mean±SD and analyzed by Mann-Whitey test. * *p*<0.05.

### Canonical PAR4 signaling contributes to venous thrombosis

With mice lacking platelet PAR4, we reported that platelet PAR4 contributes to thrombus development in the inferior vena cava stenosis model^11^. Here, we used the femoral vein electrolytic injury model to investigate venous thrombosis in real-time in mice. PAR4^R59A^ mice exhibited significantly reduced accumulation of platelets at the site of the electrolytic injury compared to control mice (**Fig. 4A+B, E**). Interestingly, accumulation of fibrin at the injury site was comparable between the control and PAR4^R59A^ mice (**Fig. 4C-E**). The data shows that canonical PAR4 activation is needed for platelet-rich thrombus formation after electrolytic injury of the femoral vein.

**Figure 4:**
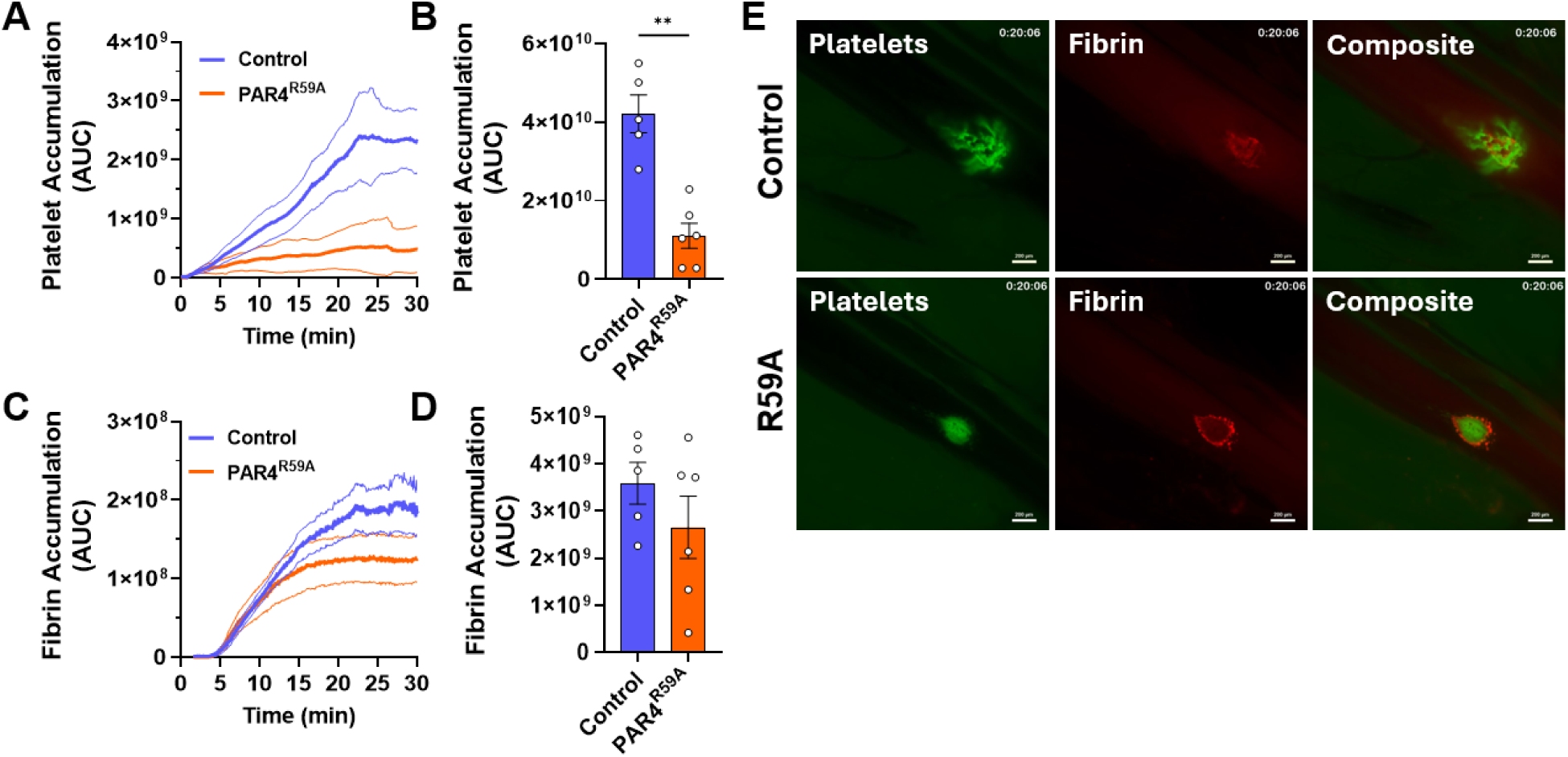
PAR4^R59A^ mice exhibit reduced platelet accumulation in the electrolytic femoral vein injury model. PAR4^R59A^ and control mice were subjected to the femoral vein electrolytic injury. Platelets were visualized with an Alexa Fluor 488-labelled anti-GPIX antibody and fibrin with an Alexa Fluor 647-labelled anti-fibrin antibody. A, B) Intravital microscopy quantification of platelet accumulation and C, D) fibrin deposition up to 30 min after injury in PAR4^R59A^ and control mice. E) Still frames from intravital microscopy videos at ∼ 20 min showing platelet accumulation (green) and fibrin generation (red) at the site of injury. Scale bar = 200 µm. Data are shown as mean±SD and analyzed by Mann-Whitney test (B, D). \*\**p*<0.01.

## Discussion

Here, we report the generation of a novel transgenic mouse carrying an Arg59 to Ala59 mutation in the canonical activation site of murine PAR4. This R59A mutation renders murine PAR4 insensitive to thrombin activation. Compared to wild-type platelets, platelets of PAR4^R59A^ mice do not aggregate after thrombin stimulation but maintain their response to PAR4AP stimulation. PAR4^R59A^ and wild-type platelets show comparable integrin activation and α-granular release after stimulation with ADP, Cvx and PAR4AP. The comparable responses to PAR4AP of wild-type and PAR4^R59A^ platelets verify that the receptor is still functional in PAR4^R59A^ mice. *In vivo*, PAR4^R59A^ mice exhibit impaired hemostasis and were protected in arterial and venous thrombosis models. The new PAR4^R59A^ mouse strain can become a useful tool to investigate loss of canonical PAR4 responses *in vivo*.

Until now analysis of global PAR4 deficiency was possibly using the transgenic PAR4^-/-^ mice^12^ or with PAR4^fl/fl^ mice^11^ after a cross with an ubiquitously expressing Cre mouse strain. Importantly, lack of PAR4 itself might lead to compensatory responses and could skew signaling pathways since PAR4 is not available to interact with its natural binding partners including P2y12^16^. The generation of the PAR4^R59A^ mice rectifies this potential complication.

Using the novel PAR4^R59A^ mouse line, we verified that thrombin-mediated activation of PAR4 has an important role in arterial and venous thrombosis and has some contribution to hemostasis in mice. The PAR4^R59A^ mutation caused impaired hemostasis with reduced platelet accumulation and fibrin deposition in a jugular vein needle puncture injury model. Together, this resulted in a prolonged bleeding time in PAR4^R59A^ compared to wild-type mice.

Our study underscores PAR4’s role in thrombosis. PAR4^R59A^ mice did not show occlusion in the carotid artery thrombosis model and had reduced platelet accumulation in the femoral vein electrolytic injury model. The dramatic antithrombotic effect in PAR4^R59A^ mice emphasize that cleavage of PAR4 at Arg59 in mice and in extrapolation at Arg47 in humans is indeed the major activation site which verifies previous findings from *in vitro* studies^5, 8, 17, 18^. Moreover, our findings with PAR4^R59A^ mice support the previous notion that thrombin-dependent platelet activation is important for thrombosis^19^.

The role of PAR4 activation in fibrin generation is not fully understood. Previous studies suggested that PAR4 activity has an impact on the generation of procoagulant platelets^20, 21^. This suggests that PAR4 activated platelets may contribute to the amplification of the coagulation cascade and subsequently fibrin generation^22^. To become procoagulant, platelets need to experience two signals such as thrombin and GPVI engagement, thrombin alone is a rather weak inducer of procoagulant platelets^23^. The simultaneous occurrence of these two signals is likely only under low shear conditions, specifically during severe vascular injury that exposes extravascular tissue factor (TF) and subendothelial collagen to the blood. In line with this, we found a PAR4-dependent role in hemostatic plug stability in a saphenous vein laser injury model^11^. One could speculate that fibrin generation becomes PAR4-dependent after a severe enough outside-in penetrating vessel injury, such as the jugular vein needle puncture injury model and not during a superficial intraluminal vessel wall activation that occurs with the electrolytic injury model. The latter injury does not lead to significant vessel wall compromise and therefore does not expose enough extracellular matrix. In line with this, a previous study found that transgenic PAR4^-/-^ mice and control mice had similar fibrin generation in the cremaster arteriole laser injury model^24^. It was suggested that thrombosis in the arterial circulation is driven by vessel injury and vessel wall TF whereas venous thrombosis is driven by so called blood-borne TF^25–27^. Under normal conditions, the levels of functional blood-borne TF is low^28^.

PAR4 was identified as potential druggable target to limit thrombotic complications since its activation needs higher thrombin concentration. These PAR4 activating concentrations are thought to occur during a thrombotic reaction^29^. The used hemostasis and thrombosis models were not able not reveal any potential non-canonical murine PAR4 signaling. It was recently suggested that human CatG can induce non-canonical signaling after cleavage of human PAR4 at serine 67 (Ser67, S67)^7, 8^. This would correspond to a cleavage at S79 in murine PAR4. Of note, murine CatG has only chymotrypsin-like activity and shows no significant cleavage activity after serine^30^. The possibility of murine PAR4 to exhibit any relevant non-canonical signaling *in vivo* is not known. Non-canonical PAR4 signaling may only occur under inflammatory conditions or in the extravascular space where thrombin is not readily generated and available. Potential cell types exhibiting non-canonical signaling under specific thrombin low conditions are, in addition to platelets, endothelial cells, smooth muscle cells, certain fibroblast populations, leukocytes including neutrophils, cardiomyocytes, epithelial cells and neurons.

Different PAR4 inhibitors were developed and tested over the time^31^. Unfortunately, the use of the experimental compounds in mice is limited due to their bioavailability, half-life and species differences. The new PAR4^R59A^ mouse can be seen as a genetic model of a systemic PAR4 inhibition *in vivo*. In addition, the mouse could be used to test the specificity and potential off-target effects of available and future PAR4 inhibitors. To reveal any significant non-canonical PAR4 signaling *in vivo*, results of PAR4^R59A^ mice should be directly compared with global PAR4 deficiency preferably using PAR4^fl/fl^ mice^11^.

In conclusion, the new PAR4^R59A^ mouse is an improved tool to analyze lack of canonical PAR4 signaling *in vivo*. The PAR4^R59A^ mice can aid the investigation of PAR4-dependent signaling in murine models of important human pathologies.

## Acknowledgements

We thank the staff of the UNC Animal Models Core for their work in generating the PAR4^R59A^ mice. We want to thank Ying Zhang for technical assistance.

